# Prepubertal gonadectomy reveals sex differences in approach-avoidance behavior in adult mice

**DOI:** 10.1101/638916

**Authors:** Kristen Delevich, Christopher Hall, Josiah R. Boivin, David Piekarski, Yuting Zhang, Linda Wilbrecht

## Abstract

Adolescence is a developmental period that is associated with physical, cognitive, and affective maturation and a time when sex biases in multiple psychiatric diseases emerge. While puberty onset marks the initiation of adolescence, it is unclear whether the pubertal rise in gonadal hormones generates sex differences in approach-avoidance behaviors that may impact psychiatric vulnerability. To examine the influence of peripubertal gonadal hormone exposure on adult behavior, we removed the gonads or performed sham surgery in male and female mice just prior to puberty onset and assessed performance in an odor-guided foraging task and anxiety-related behaviors in adulthood. We observed no significant sex differences in foraging or anxiety-related behaviors between intact adult male and female mice but found significant differences between adult male and female mice that had been gonadectomized (GDX) prior to puberty. GDX males failed to acquire the odor-guided foraging task, showed reduced locomotion, and exhibited increased anxiety-like behavior, while GDX females showed the opposite pattern of behavior. These data suggest that similar approach-avoidance phenotypes are achieved in male and female mice via different mechanisms mediated by the sex-specific hormonal milieus during pubertal maturation.

## Introduction

Across species, puberty initiates a cascade of physiological processes that induce sexual maturation and the emergence of sex differences in brain and behavior [1–3]. Adolescent changes in affective and cognitive domains may be coordinated to mediate the transition to independence [4–7]. In particular, puberty-dependent changes in the relative drive to approach novel environments versus avoid potential threats, described as ‘approach-avoidance conflict,’ may shift to promote dispersal from the natal environment and attainment of adult roles [8,9]. Approach-avoidance conflict is inherent to a wide-range of behaviors, such as a mouse foraging for food or a teenager finding a partner for a school project, and in both cases excessive approach or avoidance behavior is maladaptive [10]. Understanding the influence of puberty on approach-avoidance behavior will provide insight into normative development as well as potential mechanisms that underlie the enhanced psychiatric vulnerability post-puberty [11,12].

Many psychiatric diseases show sex differences that often emerge during adolescence [13–15]. Notably, anxiety-related disorders are more common in females than males after puberty, and in girls, earlier pubertal timing has been associated with worse mental health outcomes [13,16–21]. Several studies in humans have linked developmental changes in affective behavior and cognitive task performance to pubertal status and/or gonadal hormone concentration [22–26]. However, because adolescence is also period of increased independence and shifting social roles [27], it is difficult to disentangle the relative contributions of gonadal hormones from psychosocial changes that are secondary to pubertal changes [18,28].

Here we turned to manipulation of gonadal hormones in mouse models to understand how the rise in gonadal hormones at puberty influences approach-avoidance behavior in the context of a cognitively-demanding foraging task and anxiety-related behaviors in adult mice. We compared both male and female mice to determine whether puberty exerts different effects in males and females. In mice, as in humans, gonadal hormone production and release increases at the time of puberty onset after a quiescent period that begins shortly after birth. Increases in pulsatile release of gonadotropin releasing hormone (GnRH) from the hypothalamus stimulate pituitary gonadotropin release, which in turn promotes gonadal maturation and hormone production, increasing levels of testosterone, estradiol, and progesterone. In animal models, gonadal hormones can be manipulated via surgical removal of the gonads, or gonadectomy. By performing gonadectomy prior to the onset of puberty, we can causally test the relationship between peripubertal gonadal hormone exposure and adult behaviors [29–32].

In the current study, we sought to determine whether gonadal hormones at puberty influence adult performance in tasks that generate approach-avoidance conflict. To this end, we performed sham surgeries or gonadectomies on male and female C57Bl/6 mice before the onset of puberty and trained them as adults in an odor-guided foraging task followed by testing in the elevated plus maze (EPM) and open field to assess locomotion and anxiety-related behavior.

To interpret our data we considered three possible outcomes: A) If the sex-specific hormonal milieus at peripuberty *generate* differences in adult approach-avoidance behaviors, we would predict that adult males and females that underwent puberty will exhibit greater behavioral differences as adults than prepubertally gonadectomized mice that did not undergo puberty; B) Alternatively, it has also been shown that the sex-specific hormonal milieu can act on sexually differentiated brain circuits to *minimize* phenotypic differences between males and females [33], suggesting that prepubertal gonadectomy may result in sex differences that would otherwise be masked by peripubertal gonadal hormone exposure; C) We also considered the possibility that gonadal hormones at puberty do not play a significant role in either male or female development in our selected tasks, and in this case then outcomes should be comparable between mice that underwent prepubertal gonadectomy and same sex sham controls.

## Materials & Methods

### Animals

Male and female C57BL/6NCR mice (Charles River) were bred in-house. All mice were weaned on postnatal day (P)21 and housed in groups of 2–3 same-sex siblings on a 12:12 hr reversed light:dark cycle (lights on at 2200 h). All procedures were approved by the Animal Care and Use Committee of the University of California, Berkeley and conformed to principles outlined by the NIH Guide for the Care and Use of Laboratory Animals.

### Gonadectomy

To eliminate gonadal hormone exposure during and after puberty, gonadectomies were performed before puberty onset at P25. Mice were injected with 0.05 mg/kg buprenorphine and 10 mg/kg meloxicam subcutaneously before surgery and were anesthetized with 1–2% isoflurane during surgery. The incision area was shaved and scrubbed with ethanol and betadine. Ophthalmic ointment was placed over the eyes to prevent drying. A 1 cm incision was made with a scalpel in the lower abdomen across the midline to access the abdominal cavity. The ovaries or testes were clamped off from the uterine horn or spermatic cord, respectively, with locking forceps and ligated with sterile sutures. After ligation, the gonads were excised with a scalpel. The muscle and skin layers were sutured, and wound clips were placed over the incision for 7–10 days to allow the incision to heal. An additional injection of 10 mg/kg meloxicam was given 12–24 h after surgery. Sham control surgeries were identical to gonadectomies except that the gonads were simply visualized and were not clamped, ligated, or excised. Mice were allowed to recover on a heating pad until ambulatory and were post-surgically monitored for 7–10 days to check for normal weight gain and signs of discomfort/distress. Mice were co-housed with 1-2 siblings who received the same surgical treatment. To confirm the success of prepubertal ovariectomies, necropsy was performed on a subset of adult sham and ovariectomized mice to confirm that the uteri of OVX mice were underdeveloped compared to age-matched sham females (data not shown).

### 4 choice odor-guided foraging task

Sham or gonadectomized mice were tested in an odor-guided foraging task as young adults (P60-P70). The 4 choice odor-guided foraging task used has previously been described in detail [5,34], and within this task only the odor cue is predictive, and spatial or egocentric information is irrelevant. This task holds ethological relevance because mice use odor information to locate food sources [35]. Briefly, mice were food restricted to ~85% bodyweight by the discrimination phase. On day 1, mice were habituated to the testing arena, on day 2 were taught to dig for cheerio reward in a pot filled with unscented wood shavings, on day 3 underwent a 4 choice odor discrimination, and finally on day 4 were tested on recall of the previously learned odor discrimination immediately followed by a reversal phase. During the discrimination phase of the task, mice learned to discriminate among four pots with different scented wood shavings (anise, clove, litsea and thyme). All four pots were sham-baited with cheerio (under wire mesh at bottom) but only one pot was rewarded (anise). The pots of scented shavings were placed in each corner of an acrylic arena (12”, 12”, 9”) divided into four quadrants. Mice were placed in a cylinder in the center of the arena, and a trial started when the cylinder was lifted. Mice were then free to explore the arena and indicate their choice by making a bi-manual dig in one of the four pots of wood shavings. The cylinder was lowered as soon as a choice was made. If the choice was incorrect, the trial was terminated and the mouse was gently encouraged back into the start cylinder. Trials in which no choice was made within 3 minutes were considered omissions. If mice omitted for two consecutive trials, they received a reminder: a baited pot of unscented wood shavings was placed in the center cylinder and mice dug for the “free” reward. Mice were disqualified if they committed four pairs of omissions. The location of the four odor scented pots was shuffled on each trial, and criterion was met when the mouse completed 8 out of 10 consecutive trials correctly. 24 hours after completing discrimination, mice were tested for recall of the initial odor discrimination to criterion and immediately after reaching criterion proceeded to the reversal phase in which the previously rewarded odor (anise) was no longer rewarded, and a previously unrewarded odor (clove) now became rewarded. During the reversal phase, Odor 4 (thyme) was replaced by a novel odor (eucalyptus) that was unrewarded.

### Anxiety testing

Mice underwent anxiety testing >1 week after completing the odor-guided reversal task and returned to ad libitum feeding. All testing took place during the last 1-2 hours of the light phase (0800-1000 h) and mice were allowed to habituate to the testing room in their home cages for 30 minutes prior to testing. Mice were sequentially run in the elevated plus maze immediately followed by the open field test.

### Elevated plus maze

The mouse was placed in the center of an elevated plus maze and allowed to explore freely for 10 min. The EPM was made out of opaque white acrylic consisting of 2 open arms (30 cm long by 6 cm wide), 2 closed arms (30 cm long by 6 cm wide, with 20.5 cm high walls on the sides and end of each arm), and a center square (6 cm by 6 cm). The closed arms of the EPM were attached to a stable platform raised 66 cm from the floor. The total time spent in each zone (center, closed, open) was analyzed using EthoVision software (Noldus; Sacramento, CA). The chamber was cleaned with 70% ethanol and allowed to dry between tests of mice. The EPM was performed with room lights on (260 lx).

### Open field

Immediately after completing EPM testing, the mouse was transferred to a clear acrylic open field arena (42 cm by 42 cm floor dimensions, with four walls that were each 30.5 cm high, and no ceiling) for 15 min. The acrylic open field arena was located inside a sound-attenuated chamber (Med Associates; Fairfax, VT) with lights on (40 lx inside the chamber). Locomotion was monitored using infrared beam breaks (Versamax, AccuScan Instruments; Columbus, OH). Total distance covered and percent of time spent in the center (defined as > 7.875 cm from the edges of the chamber, i.e. 3 grid squares in Versamax analysis software), as well as vertical rears were analyzed. Data suggest that unsupported vertical rears are an exploratory movement that is sensitive to environmental context and stressors [36]. As described in Sturman et al., we quantified supported and unsupported rears based on the zone in which they occurred (outer zone vs. center zone, respectively). The chamber was cleaned with 70% ethanol and allowed to dry between testing of individual mice. Mice were returned to the home cage immediately after the open field test.

### Statistical analysis

For comparisons between 2 groups, a t-test was used when data were normally distributed, and Welch’s correction was applied when variance was unequal. The D’Agostino & Pearson test was used to test for normality. Groups that were not normally distributed were compared using a Mann Whitney U test. When 4 sex by surgery groups were considered simultaneously and data was normally distributed, a 2-way ANOVA was performed with sex and surgery as factors. For experiments in which only 3 groups were compared, a one-way ANOVA was performed. Post-hoc comparisons were then performed as described above (i.e. using a t-test or Mann Whitney U test). For the analysis shown in Fig. 2 and Figs. S1-3, planned comparisons were performed to address two *a priori* questions: 1) Do adult intact males and females differ in behavior? 2) Does gonadectomy alter performance compared to intact mice of the same sex? As these were separate *a priori* questions, multiple comparisons corrections were not applied to these tests.

## Results

### Prepubertal castration but not ovariectomy reduces likelihood of completing 4 choice odor-guided foraging task

We performed sham surgery or gonadectomy (GDX) on male and female mice at P25, generating four behavioral groups: intact males, intact females, castrated (Cast) males, and ovariectomized (OVX) females that were trained in an odor-guided foraging task (between P60-70) followed by open field and elevated plus maze testing (between P80-90) (Figure 1A). At the initiation of training, we compared body weights across groups and found a main effect of sex [F(1,63)= 86.37, p<0.0001] but no main effect of surgery [F(1,63)= 2.487, p= 0.1198] (Figure 1B). To motivate performance in the odor-guided foraging task, mice were food restricted to ~85% of their initial body weight and trained across four days, including 1) habituation, 2) shaping, 3) discrimination, and 4) recall and reversal task phases (Figure 1C, see methods for more detail). Over the course of training, we found that only (1/13) Cast males successfully completed the reversal phase of the task, with (11/13) mice dropping out before or during the discrimination phase (Figure 1D). While there was no significant difference in the proportion of intact males (13/18) vs. intact females (10/15) that completed the task (p > 0.99 Fisher’s exact test), there was a significant difference in the proportion of Cast males (1/13) vs. OVX females (14/15) that completed the task (p<0.0001, Fisher’s exact test) (Figure 1D). The proportion of Cast males that completed the task was also significantly lower compared to intact males (p= 0.0007, Fisher’s exact test), whereas OVX and intact females did not significantly differ (p= 0.17, Fisher’s exact test). Dropout among Cast males was driven by 1) a failure to complete the shaping phase within the allotted 3 hours (3/13) or 2) disqualification during the discrimination phase due to omission trials (4 pairs of omissions) (9/13) (Figure 1D). There was a significant interaction between sex and surgery on the number of omissions made during discrimination [F(1, 56) = 9.33, p= 0.0034]. Cast males made significantly more omissions compared to intact males (p= 0.0018, Tukey’s multiple comparisons test) and OVX females (p<0.0001, Tukey’s multiple comparisons test) (Figure 1E). Meanwhile, intact males and females did not differ from each other in the number of omissions made during discrimination testing (p= 0.36, Tukey’s multiple comparisons test) (Figure 1E). We next asked whether differences in body weight loss explained drop out among Cast males, reasoning that diminished weight loss could contribute to lower task motivation. However, we found no significant effect of surgery on percent initial body weight at discrimination phase [F(1,53) = 0.065, p= 0.80] (Figure 1F) indicating that Cast males lost a similar percentage of their body weight compared to other groups. Therefore, we were unable to further examine 4 choice odor-guided foraging task behavior in Cast males, including reversal learning.

**Figure 1.**
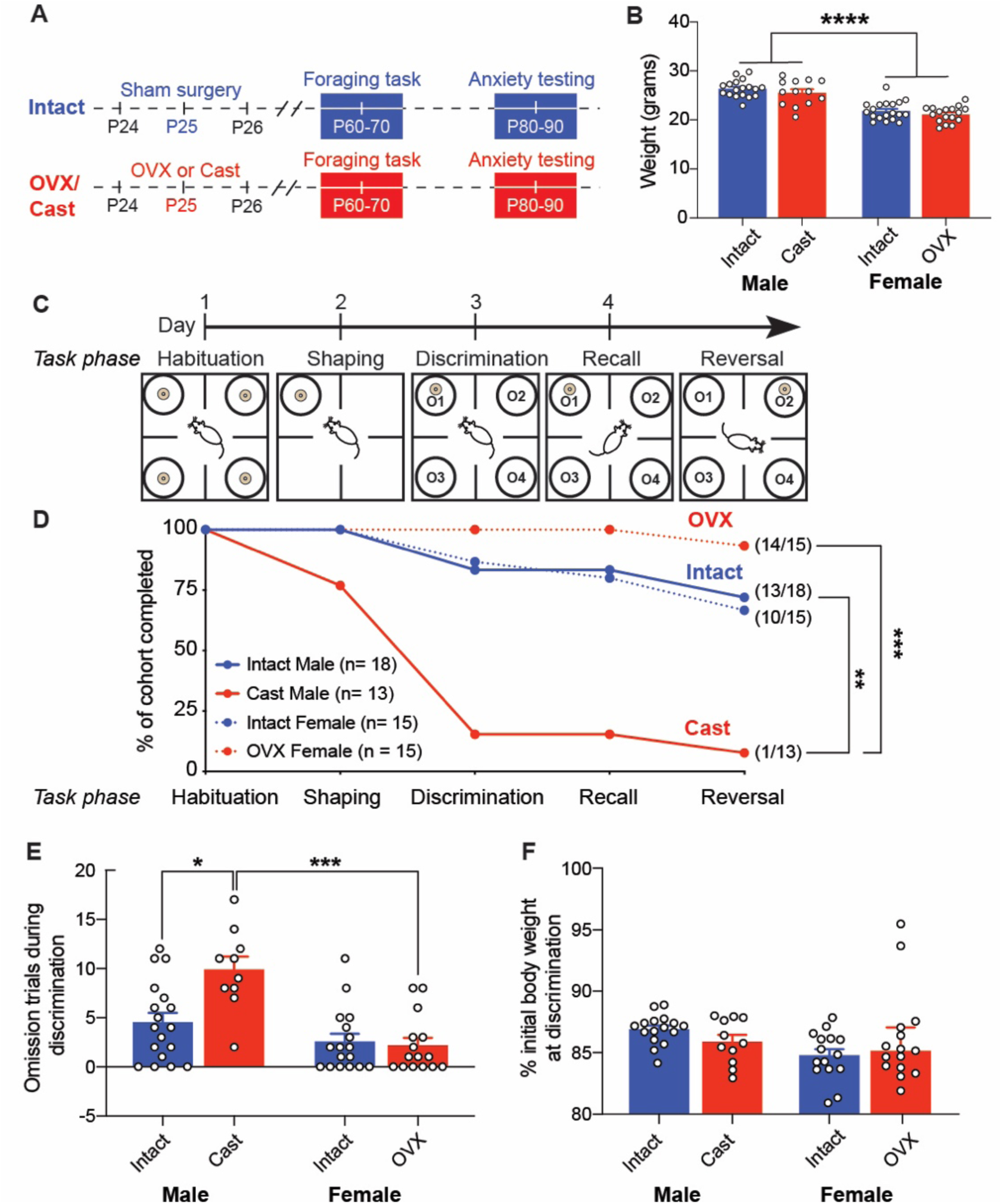
Prepubertal castration but not ovariectomy reduces likelihood of completing foraging task. A. Male and female mice were weaned at postnatal day (P)21 and underwent sham or castration/ovariectomy at P25 before the onset of puberty. Mice were then trained in a 4 choice odor-guided foraging task in early adulthood (P60-P70) followed by anxiety testing from P80-P90. B. There was a significant main effect of sex on initial weight before food restriction began [F(1, 63) = 86.37, p<0.0001] but no significant effect of surgery (sham vs. gonadectomy) on initial weight [F(1, 63) = 2.487, p = 0.1198]. C. Overview of 4 choice odor-guided foraging task. D. Completion rate of the four cohorts: Intact Male, Intact Female, castrated (Cast) Male, and ovariectomized (OVX) Female across the four days of behavioral training (aligned to panel C). There was a significant difference in the completion rate at reversal phase between Cast males and OVX females (p<0.001, Fisher’s exact test with Bonferroni correction) but no significant difference in completion rates between intact males and females (p>0.99, Fisher’s exact test with Bonferroni correction). Cast males also significantly differed from intact males (p<0.01, Fisher’s exact test with Bonferroni correction). E. Dropout by Cast males was largely driven by omission trials (no choice made within 3 min trial) during the discrimination phase. F. Weight loss at discrimination phase. Neither gonadectomized group (Cast or OVX) significantly differed from intact groups, suggesting that differences in weight loss did not account for drop out by Cast males. All data presented as mean ± SEM.

### Prepubertal ovariectomy reduces bias for perseverative errors during reversal compared to intact females

We next examined performance in the odor-guided foraging task across intact males, intact females, and OVX females. During the discrimination phase (Figure 2A), there was no significant difference among groups in trials to criterion (p= 0.85, Kruskal Wallis ANOVA) (Figure 2B) or total errors (p = 0.85, Kruskal Wallis ANOVA) (Figure 2C). The following day mice were tested for their recall of discrimination learning, and groups did not differ in performance (**Figure S1**). During a reversal phase that immediately followed the completion of the recall phase (Figure 2D), there was no significant difference in trials to criterion (p= 0.52, Kruskal Wallis ANOVA) (Figure 2E) or total errors (p= 0.28, Kruskal Wallis ANOVA) (Figure 2F) across groups. Next, we more closely examined the types of errors that mice made during reversal. All groups showed a strong bias for selecting the odor that was previously rewarded in the discrimination phase (**Figure S1**). We divided errors to the previously rewarded odor into 2 subtypes: perseverative (errors made before the first correct trial) and regressive (errors made after first correct trial) errors. Perseverative errors reflect a tendency to stick to a previously learned rule, while regressive errors reflect a failure to acquire or maintain the new rule. We found that groups did not significantly differ in the number of perseverative errors made (p= 0.60, Kruskal Wallis ANOVA), but OVX females made significantly more regressive errors than intact females (U= 36, p= 0.0459 uncorrected) (**Figure S1**). We next examined the pattern of perseverative and regressive errors made by individual mice. While we did not find a difference in the pattern of reversal errors between intact males and females (U= 71.5, p= 0.3778), we found that intact females exhibited a significantly higher bias for perseverative vs. regressive errors compared to OVX females (intact females vs. OVX females: U= 47.5, uncorrected p= 0.0375) (Figure 2G). In addition, we observed that intact females accumulated rewards at a significantly higher rate after the first rewarded trial in reversal compared to OVX females (Figure 2H).

**Figure 2.**
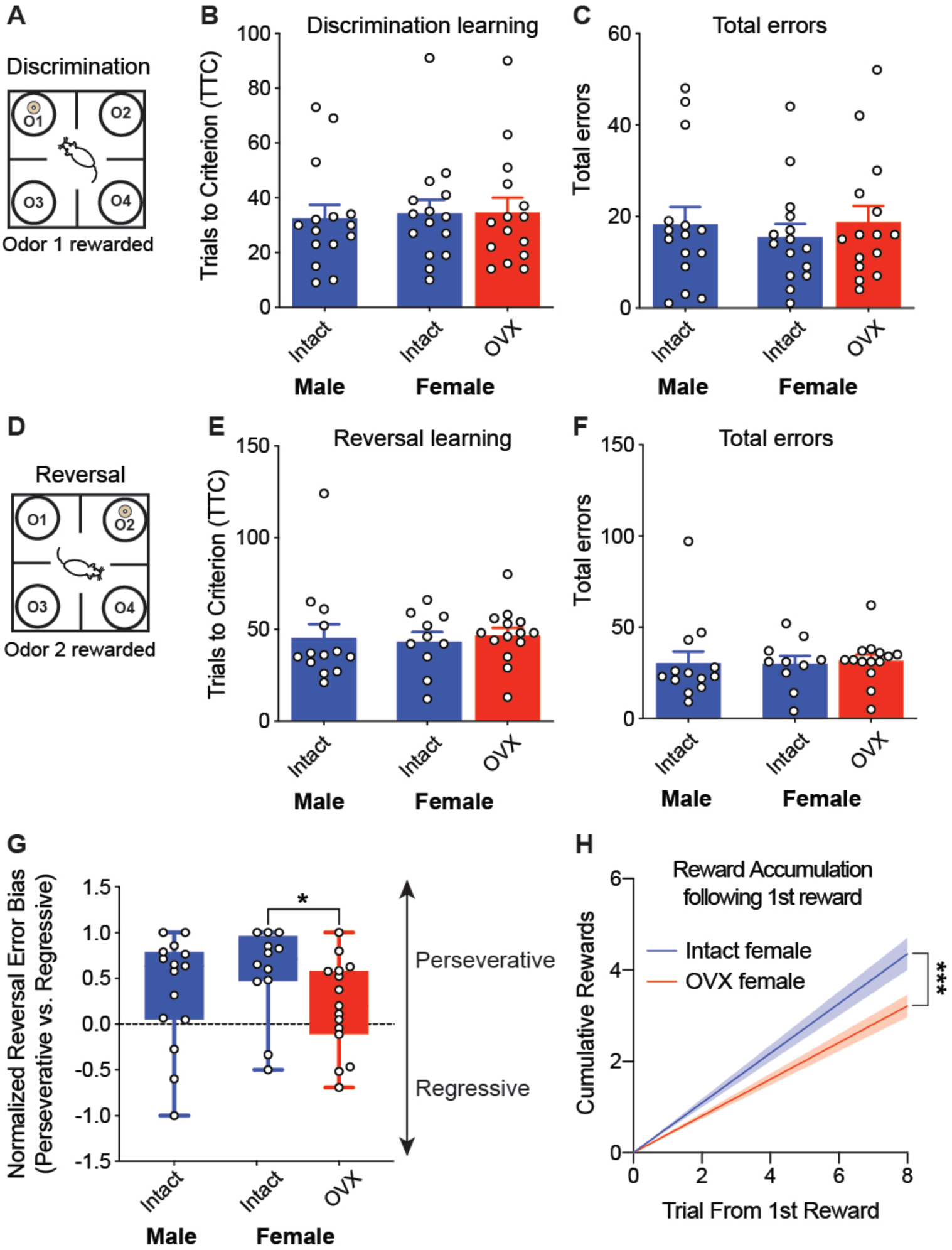
Prepubertal ovariectomy reduces bias for perseverative errors following reversal compared to intact females. A. Overview of discrimination phase when mice learn that Odor 1 is rewarded. B. All cohorts exhibit similar trials to criterion (TTC) to complete discrimination. C. All cohorts made similar number of errors during the discrimination phase. D. Overview of reversal phase when mice learn that Odor 1 is no longer rewarded, and now Odor 2 is rewarded. E. All cohorts exhibited similar trials to criterion (TTC) to complete reversal. F. All cohorts exhibited similar total number of errors during the reversal phase. G. Intact female mice exhibited greater bias towards perseverative vs. regressive errors compared to OVX females (p= 0.0375, uncorrected Mann-Whitney U test). Error bias was calculated for each mouse as (perseverative – regressive errors) / (perseverative + regressive errors). H. After the first rewarded trial in reversal, intact females accumulated rewards at a higher rate than OVX females (p< 0.0001, extra sum-of-squares F test).

### Prepubertal gonadectomy has opposing effects on anxiety-related behavior in approach-avoidance conflict tests in adult males and females

To determine whether prepubertal gonadectomy affects ‘anxiety-like’ behavior in tasks that rely on approach-avoidance conflict, we tested mice from P80-90 in the elevated plus maze (EPM) and open field [37]. In the elevated plus maze (Figure 3A), there was a significant interaction between sex and surgery on locomotion [F(1,53) = 14.84, p= 0.0003], such that differences in total distance travelled emerged between Cast males and OVX females (p<0.0001, Tukey’s multiple comparisons test) that were not present between intact males and females. Cast males traveled significantly less distance compared to OVX females (Figure 3B, **Figure S2**). In addition, there was a significant interaction between sex and surgery on time spent in the closed arms of the EPM [F(1, 53) = 10.67, p<0.01], with Cast males spending significantly more time in the closed arms compared to OVX females (p<0.001, Tukey’s multiple comparisons test), while intact males and females did not differ from each other (p= 0.99, Tukey’s multiple comparisons test) (Figure 3C,D, **Figure S2**). Immediately after completing EPM testing, mice were run in the open field test (Figure 3E). Similar to the EPM results, there was a significant interaction between sex and surgery on locomotion, and again, OVX females traveled greater distance compared to Cast males (Figure 3F, **Figure S2**). There was a significant interaction between sex and surgery on time spent in the outer zone of the open field [F(1, 58) = 18.56, p<0.0001], with Cast males exhibiting more anxiety-like behavior (time in outer zone) compared to OVX females (p<0.001 Tukey’s multiple comparisons test) (Figure 3G, **Figure S2**). In addition, Cast males made significantly fewer vertical rears in the open field compared to OVX females (p< 0.0001, Tukey’s multiple comparisons test). We confirmed that across all animals, time spent in the closed arms of the EPM and time spent in the outer zone of the open field were significantly correlated (**Figure S2**). Finally, studies suggest that unsupported vertical movement, or rearing, is an exploratory movement that is sensitive to environmental context and stressors [36]. We found that Cast males made fewer unsupported rears compared to intact males (U= 30, p= 0.0003) but there was no significant difference between intact females and OVX females (**Figure S3**).

**Figure 3.**
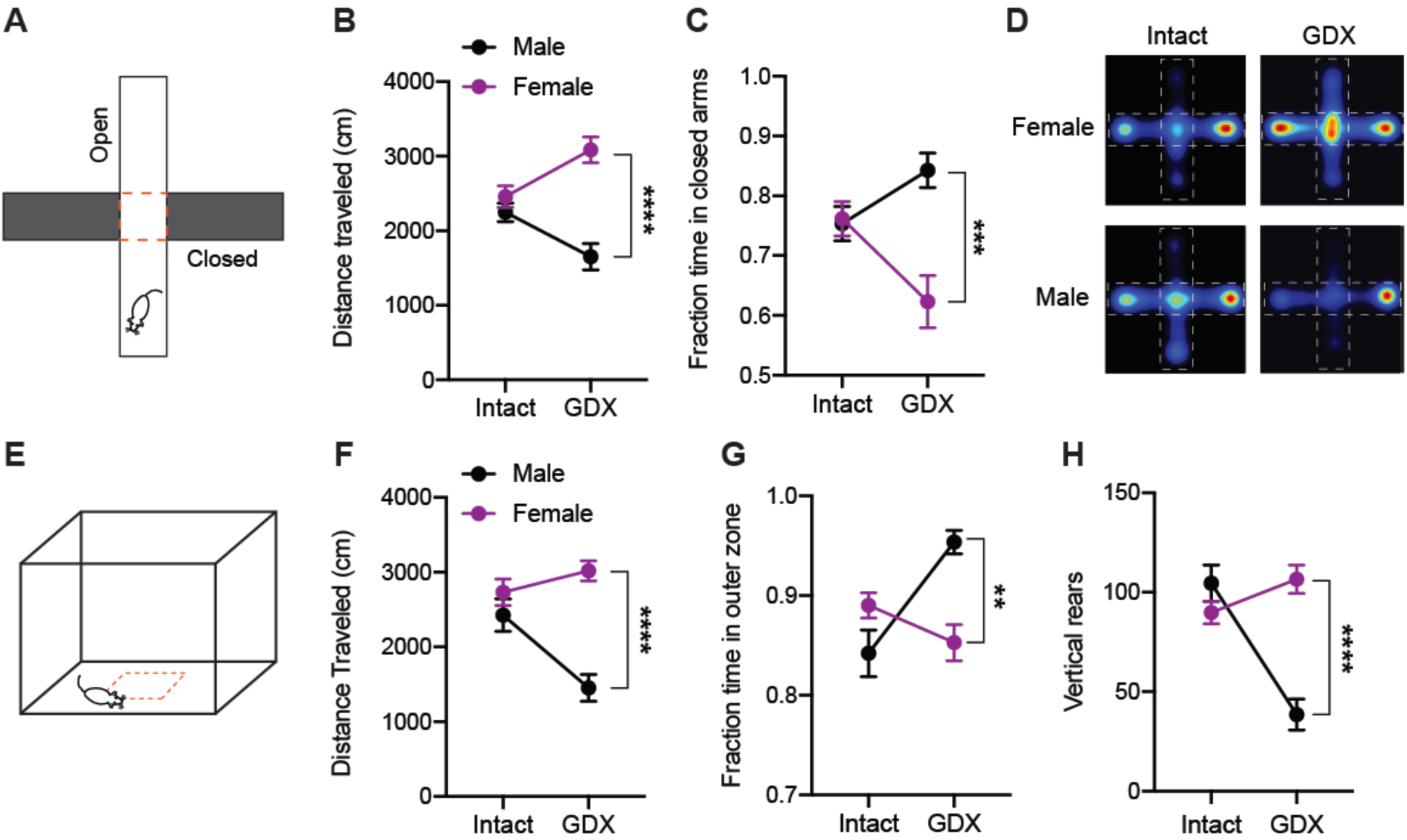
Prepubertal gonadectomy produces sex differences in adult anxiety-related behavior not present in intact males and females. A. Elevated plus maze (EPM) apparatus. B. There was a significant interaction between sex and surgery on total distance travelled during the EPM test [F(1,53)= 14.84, p= 0.0003]. C. There was a significant interaction between sex and surgery on fraction of time spent in the closed arms during the EPM test [F(1, 53)= 10.67, p= 0.0019]. D. Representative heat maps showing time spent in open and closed arms for Intact female, OVX female, Intact male, and Cast male groups. E. Open field. F. There was a significant interaction between sex and surgery on total distance travelled during the open field test [F(1, 58)= 11.89, p= 0.0011]. G. There was a significant interaction between sex and surgery on time spent in the perimeter (not in center) of the open field [F(1, 58)= 18.56, p< 0.0001]. H. There was a significant interaction between sex and surgery on vertical rears made in the open field [F(1, 58)= 30.20, p< 0.0001]. **p< 0.01, ***p< 0.001, ****p< 0.0001 Tukey’s multiple comparisons test.

### Some effects of prepubertal gonadectomy on anxiety-related behavior in approach-avoidance conflict tests are modified by age

A previous study from our lab compared behavior in the EPM and open field test between intact and prepubertally gonadectomized mice during a mid-adolescent time point (P40). Given that the mouse strain, surgical manipulation, and anxiety-related testing procedure were consistent, we compared the effect size of gonadectomy on anxiety measures at adolescent (P40-47) and adult (P80-90) time points to determine whether effects are stable or change with age. In males, we found that the effect of prepubertal castration on anxiety-like behavior was stable between P40 adolescent and P80 adult cohorts (Figure 4A,B) but the negative effect of prepubertal castration on locomotion was more pronounced in the P80 adult cohort, particularly in the open field (Cast P40 vs. Cast P80 U= 24, p= 0.0013) (Figure 4C,D). For females, however, we found a significant difference in the effect of prepubertal ovariectomy on time spent in closed arms of the EPM in adults compared to adolescent females, with OVX being associated with less time spent in closed arms in the P80 adult cohort but not the younger P40 cohort (intact P40 vs. OVX P40: t_42_= 0.3827, uncorrected p= 0.70; intact P80 vs. OVX P80: t_30_= 2.752, p= 0.01) (Figure 4E). Thus, the anxiolytic effect of OVX on female EPM behavior emerged only with age (P40 OVX vs. P80 OVX: t_27_= 3.106, p= 0.0055). Meanwhile, OVX and intact female mice did not differ in open field behavior at either age (Figure 4F). Finally, a similar pattern was found for locomotion in the EPM task, where a significant increase in distance traveled by OVX compared to intact females was observed at P80 but not P40 (intact P40 vs. OVX P40: t_42_= 1.334, uncorrected p= 0.1895; intact P80 vs. OVX P80: t_30_= 2.796, p= 0.0089), and the OVX effect differed significantly between P40 and P80 (t_18.02_= 2.43, p= 0.0257) (Figure 4G). We did not observe a significant difference between OVX and intact female for distance traveled in open field behavior at either age (Figure 4H).

**Figure 4.**
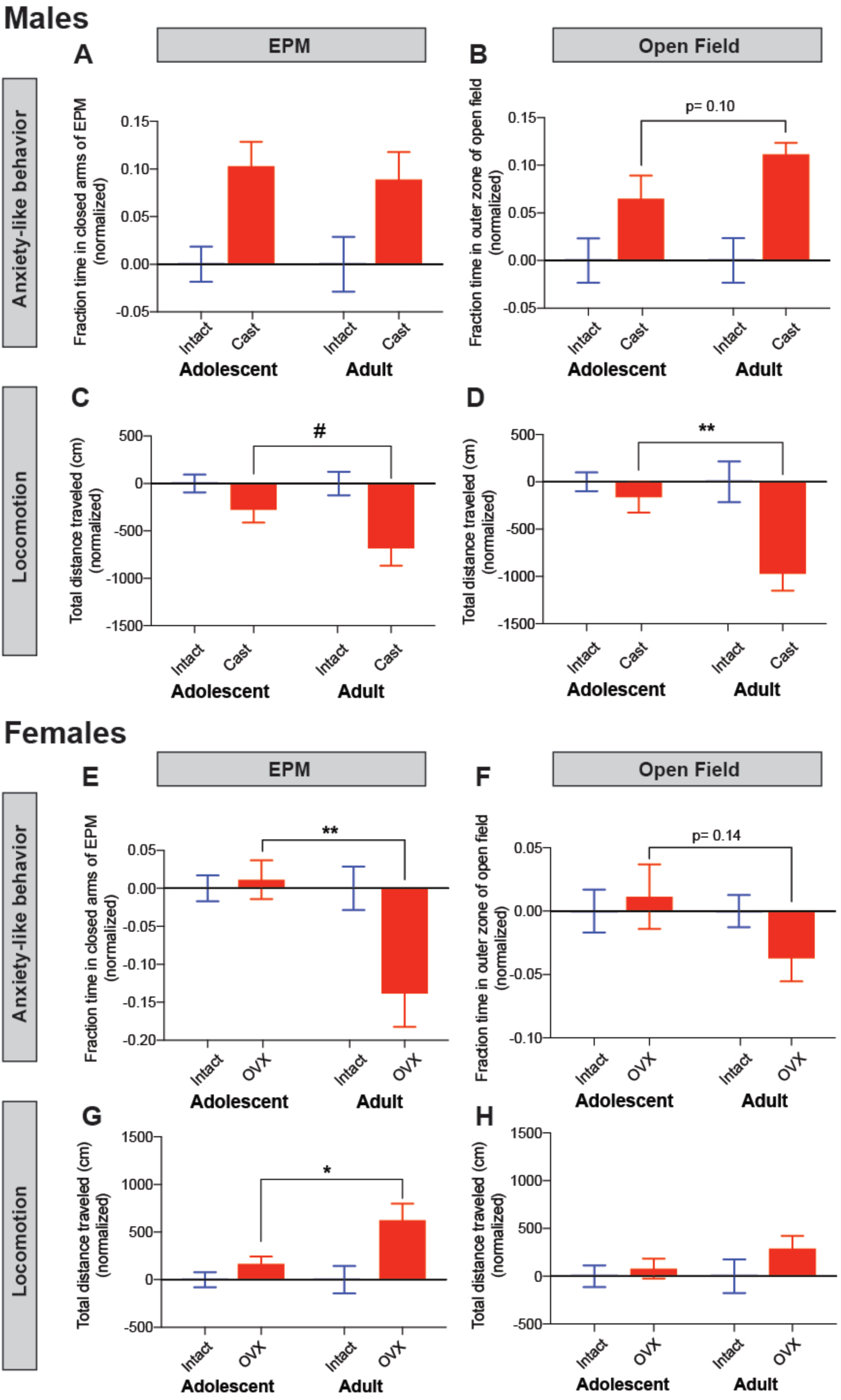
Effects of prepubertal gonadectomy on anxiety-related behavior and locomotion in approach-avoidance conflict tests are modified by age. A. Cast males spend more time in the closed arms of the EPM compared to intact males, and the magnitude of this effect does not change between P40 and P80. B. The effect of prepubertal Cast on open field anxiety-like behavior did not significantly change between P40 and P80. C. There was a trend for Cast males to move less relative to intact males in the EPM at P80 compared to P40 (t_20_= 1.849, p= 0.0793). D. Cast males traveled significantly less distance in the open field relative to intact males at P80 compared to P40 (U=24, p= 0.0013). n= 23 P40 intact males; n= 12 P40 Cast males, n= 16 P80 intact males; n= 10 P80 Cast males. E. A significant reduction in anxiety-like behavior in the EPM emerged at P80 for OVX compared to intact females that is not present at P40 (t_30_= 2.752, p= 0.01). F. The effect of prepubertal OVX on open field anxiety-like behavior did not significantly change between P40 and P80. G. OVX females traveled greater distance in the EPM relative to intact females at P80 compared to P40 (t_18.02_= 2.431, p= 0.0257). H. There was no significant effect of OVX on distance traveled in open field at either age. n= 29 P40 intact females; n= 15 P40 OVX females; n= 18 P80 intact females; n= 14 P80 OVX females. #p<0.1; *p<0.05; **p<0.01.

### Anxiety-like behavior in open field is associated with foraging task drop out in males but not females

Finally, we sought to examine whether differences in learning and performance in the odor-guided foraging task showed a relationship with EPM and open field behavior, because all three tasks to some extent task the tradeoff between approach and avoidance behavior. We noted that there was a clear parallel between the opposing effect of prepubertal gonadectomy in males vs. females on the likelihood of completing the odor-guided foraging task and measures of anxiety-like behavior: Cast males exhibited the highest measures of anxiety-like behavior and completed the foraging task at the lowest rate, while OVX females exhibited the lowest anxiety-like behavior (in EPM) and completed the foraging task at the highest rate. We first compared anxiety-related measures between mice that completed the odor-guided foraging task (“completers”) and those that dropped out due to omissions (“drop-outs”). In males, we found that the time spent in the outer zone of the open field was significantly higher in “drop-outs” compared to “completers” (p= 0.0003, Sidak’s multiple comparisons test) (Figure 5A). This effect was not purely driven by Cast males, as it was present when intact males were examined separately (**Figure S4**). One potential confound is that mice who complete the odor-guided foraging tasks experience more handling and spend more time in the foraging task arena, which shares some characteristics with the open field. This may also explain why we did not also find a difference in time spent in the closed arms of the EPM between male drop-outs and completers.

**Figure 5.**
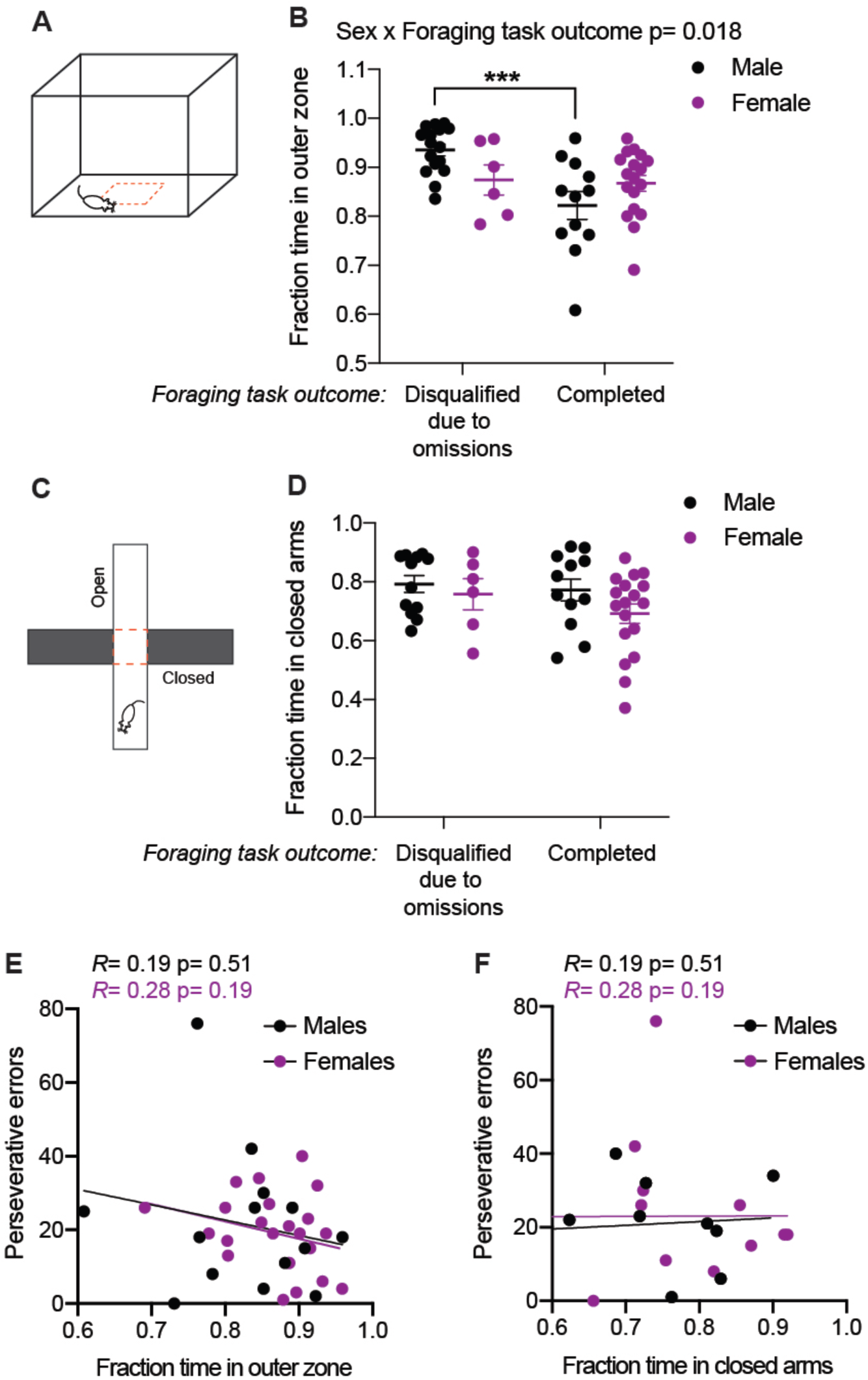
Drop out in foraging task is associated with anxiety-like behavior in the open field for males but not females. A. Open field. B. There was a significant interaction between sex and foraging task outcome (disqualified or completed) on fraction time in the outer zone [F(1, 48)= 5.98, p= 0.018]. Males who dropped out of the foraging task spent significantly more time in the outer zone of the open field compared to males who completed the task (p= 0.0003, Sidak’s multiple comparisons test). C. Elevated plus maze. D. There was no main effect of reversal task outcome on time spent in the closed arms of the EPM. E. Fraction time in the outer zone of the open field showed no significant relationship with perseverative errors in the reversal phase. E. Fraction time in the closed arms of the EPM showed no significant relationship with perseverative errors in the reversal phase of the odor-guided foraging task.

In females, we found no significant difference in anxiety-related measures between drop-outs and completers, and time spent in the closed arms of the EPM did not differ between intact male or female mice that dropped-out or completed the task (Figure 5B). There are two important caveats to this finding: 1) our statistical power to detect differences between drop-outs and completers in the female group was reduced compared to males because fewer females overall (intact + OVX) were disqualified and 2) we did not control for phase of the estrous cycle when females were tested in the foraging task vs. anxiety-related tests. Previous data have shown that anxiety-related behavior fluctuates along the estrous cycle [38], so intact females may not have been in the same state between testing conditions.

Finally, we tested whether variation in anxiety-related behavior showed a relationship with flexibility in behavioral updating following rule reversal [39,40]. We examined the relationship between measures of anxiety-related behavior collected in the EPM and open field and perseverative errors made during the reversal task. We found no significant correlation between these variables in either males or females (Figure 5C,D).

## Discussion

Peripubertal gonadal hormone exposure has long-lasting effects on sexual and social behaviors [41–46], yet far less is known about how the rise in gonadal hormones at puberty affects cognitive and affective development. Standard models of gonadal hormone activity suggest the rise in gonadal hormones at puberty could drive the emergence of sex differences in cognitive and affective domains. Alternate models suggest the rise in peripubertal gonadal hormones could also *prevent* behavioral differences from arising between males and females [33]. Also, we must consider that many aspects of development may occur at the same time as puberty but be independent of pubertal hormone changes [47,48].

In the current study, we used gonadectomy to test the role of peripubertal gonadal hormones in one cognitive and two anxiety-related tasks that generate approach-avoidance conflict. We were particularly interested in these approach-avoidance related tasks because adolescence is a period during which enhanced approach is likely adaptive for gaining independence, and previous studies have shown that exploration of novel environments increases across adolescence in male and female mice [49–51]. It has been hypothesized that pubertal gonadal hormones may alter brain function to promote dispersal from the natal environment and to facilitate behaviors such as foraging that require exploration [8,30], but see [52]. We previously showed that prepubertal gonadectomy increased anxiety-like behavior in males (but not females) during adolescence [29]. Our current data provide further support for the role of peripubertal testicular hormones in tuning approach-avoidance behavior: Cast males exhibited a lack of engagement in the foraging task and displayed reduced exploratory locomotion and increased anxiety-like behavior in the open field, all consistent with reduced approach behavior and/or greater avoidance. In boys (14-17 years old), lower testosterone levels were associated with increased anxiety and depressive symptoms [53]. It is interesting to note that juvenile males, who secrete testicular hormones at much lower levels than adults, readily acquire the odor-based foraging task and exhibit more efficient reversal learning when compared to adult males [9]. Therefore, the behavioral requirement for testicular hormones may change in an age-dependent manner.

In adulthood, we observed no major differences in locomotion or anxiety-related behavior between gonadally intact adult males and females, consistent with previous data from our lab (Boivin et al. 2017) and others [54]. Notably, we did find highly significant differences in both locomotion and anxiety-like behavior when we compared prepubertally gonadectomized males (Cast) and females (OVX) (Figure 3). These data are consistent with less well-known models of hormonal action in which gonadal hormones act via different pathways in each sex to ultimately produce more similar behavioral phenotypes (De Vries 2004). It is useful to point out that tests of anxiety-related behavior may not capture the same underlying processes in males and females; for example, it has been reported that the behavior of female rats in the EPM is driven more by locomotion than anxiety, whereas in males the reverse is true [55]. We therefore interpret that the divergent phenotypes we observed in Cast males and OVX females could be driven by different mechanisms that shift behavior along an approach-avoidance continuum [56]. Future studies should be designed to address what underlying processes drive the different behavioral phenotypes we observed.

Given the link between puberty onset and the emergence of female bias for anxiety-related disorders, we were particularly interested to examine the role of gonadal hormone exposure on approach-avoidance behaviors in females. We observed that as adults, females ovariectomized before puberty onset exhibited more exploratory behavior in the EPM (less time spent in closed arms) compared to intact females. This OVX versus sham difference was significant in adult mice (P80) but was not present when we sampled at a P40 adolescent time point, suggesting it emerged over time. In addition, our group previously showed that prepubertal ovarian hormone exposure (that advanced puberty onset) did not acutely increase anxiety-like behavior in juvenile female mice [29]. Taken together, these data suggest that peripubertal ovarian hormone exposure does not have acute anxiogenic effects but that it may have organizational or activational effects that tune approach-avoidance balance over a longer period of time. These data suggest ovarian hormones may play a role in anxiety-related behavior in females, but does not rule out the potential importance of modifiable factors such as environment and socialization (Graber 2013). The delayed emergence of the prepubertal ovariectomy effect on anxiety in females is in contrast to the timing of the prepubertal castration effect on anxiety-like behavior, which was present at the adolescent time point. This difference suggests that males and females exhibit different trajectories for the maturation of anxiety-related behaviors, and is consistent with testicular hormones having an activational effect on approach-avoidance behaviors [57].

Previous studies indicate that in some cases male and female rodents utilize different decision making strategies [58], which may reflect ethologically relevant differences in how they use information in their environment [59]. Such reported sex differences in rodents include reduced risk preference in females compared to males [60–62], greater impulsivity in females under mild food restriction [63], and slower acquisition by female rats of a visual discrimination task [64]. While we did not find significant differences in performance between intact males and females in the odor-guided foraging task, we found that intact females exhibited a stronger bias for perseverative vs. regressive errors compared to OVX females, suggesting that peripubertal ovarian hormone exposure altered foraging strategy. This bias for perseverative errors reflects a difference in the strategy employed during reversal: intact females persisted in the old rule (Odor 1= rewarded) longer, but once they chose Odor 2 and were rewarded, they acquired the new rule faster and accumulated rewards at a higher rate compared to OVX females. This pattern of behavior by intact females may reflect reproductive pressure to exploit known available food sources, whereas OVX females exhibited more exploratory foraging strategy.

Our lab has previously shown that advancing puberty in juvenile female mice alters reversal performance in the same odor-guided foraging task [30]. Pubertal advancement increased the trials to criterion during reversal compared to age-matched prepubertal mice [30], consistent with juveniles exhibiting faster reversal/greater flexibility compared to adults [5]. While the prepubertal OVX manipulation significantly altered reversal strategy in adult females, we did not observe any associated improvement in performance (i.e. a reduction in trials to criterion) suggesting that prepubertal OVX did not extend “juvenile-like” performance to P60 females. We previously showed that performance in the reversal task of the odor-guided foraging task is dependent on the dorsomedial prefrontal cortex (dmPFC) of mice [5] and that prepubertal ovariectomy has long-lasting effects on inhibitory transmission in the dmPFC [30]. Future experiments will explore mechanisms by which peripubertal gonadal hormone exposure influences approach-avoidance behavior and its relationship to decision-making [65], with late-developing circuits of particular interest [66].

Our current study has several limitations. First, we did not track the hormonal status of intact females. In part this was due to the foraging task design – because it occurs over 6 days and sessions are not repeated, we were unable to sample individual mice across the different phases of the estrous cycle. The fact that we were able to detect significant differences in behavior compared to OVX females in locomotion, anxiety-related behavior, and foraging strategy, suggests that these effects are robust to variance in ovarian hormone levels present in intact females. Next, our study design does not enable us to distinguish between activational and organizational effects of testicular and ovarian hormones. Future studies could control timing of gonadectomy and use hormone replacement to determine whether peripuberty is a sensitive period for the organization of anxiety-related behavior and foraging strategy. Regionally specific hormone replacement [67] and cell-type specific manipulation of steroid hormone receptor expression [68] could elucidate the neural mechanisms underpinning the effects of pubertal gonadal hormones on the behaviors examined here.

In rodent models, it is well known that species and strain differences, as well as environmental stressors during rearing such as shipping [69], can affect behavioral outcomes. Our current results conflict with reports in rats, where prepubertal castration reduced anxiety-like responses in adulthood [70,71] and increased ambulation [72]. More generally, sex differences in anxiety-related behavior and locomotion appear more consistently in rats, with female rats ambulating more and spending more time in anxiogenic zones compared to males [73]. However, in line with our findings, Kim and Spear observed sex-specific effects of prepubertal gonadectomy on social anxiety in rats: males displayed anxiety-like decreases in social preference whereas females displayed anxiolytic-like increases in social preference [74]. In mice, other studies have found evidence for reduced anxiety-like behavior in intact males compared to intact females [75,76], and one study did not find an effect of prepubertal castration on anxiety-like behavior in the open field [77]. Therefore, the effects of species, strain, and rearing conditions should be taken into account when interpreting sex and gonadal hormone dependent differences in approach-avoidance behavior.

## Conclusion

The current study sought to determine whether gonadal hormones at puberty influence approach-avoidance behavior in a sex-specific manner. We found that prepubertal gonadectomy in mice revealed significant differences in approach-avoidance behaviors that were not present between intact male and female mice suggesting that similar approach-avoidance phenotypes are achieved in male and female mice via different mechanisms mediated by gonadal hormones from puberty into adulthood.

## Acknowledgments

We thank Jessica Wahlberg and Kenechukwu Okwuosa for technical assistance with mouse behavior testing. We thank Dr. George Prounis and Dr. Irv Zucker for their feedback on the manuscript.

