## Supplementary Material for "Prepubertal gonadectomy reveals sex differences in approach-avoidance behavior in adult mice"

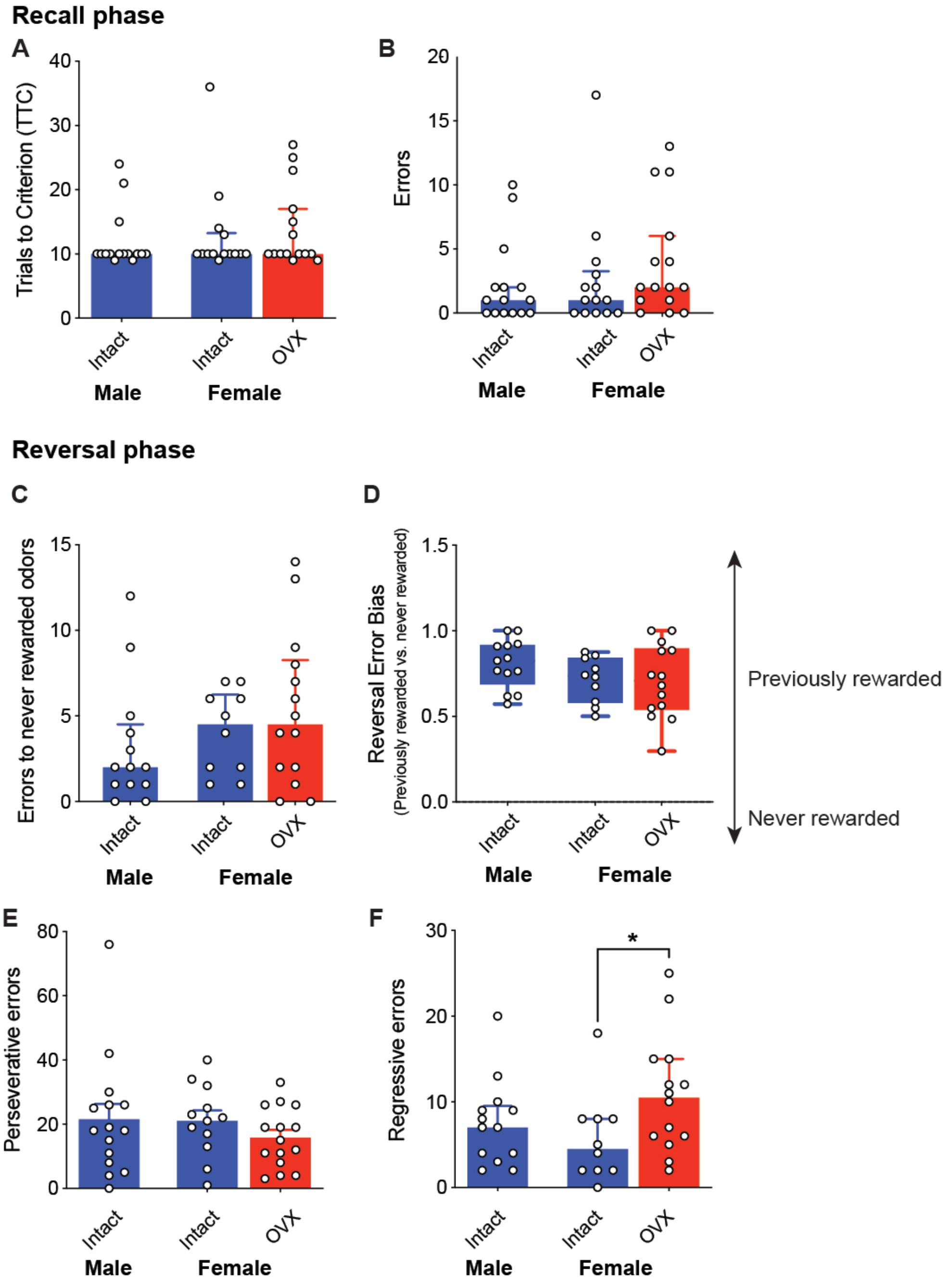


**Figure S1. OVX females significantly differ from intact females by the number of regressive errors made during reversal, but not other measures (related to Figure 2).** A. Groups did not differ in trials to criterion in the recall task phase (H= 0.89, p= 0.64). B. Groups did not differ in the number of errors made during recall (H= 2.51, p= 0.29). C. Groups did not differ in the number of errors made to odors that were never rewarded (Odor 3 and Odor 4) (H= 2.31, p= 0.31). D. There was no difference across groups in bias for previously rewarded vs. never rewarded odors (F(2, 31.12)= 1.52, p= 0.23, Brown-Forsythe ANOVA). E. Groups did not differ by total number of perseverative errors made in reversal (H= 1.3, p= 0.52). F. OVX females made more regressive errors during reversal compared to intact females (U= 36, p= 0.046 uncorrected). Data in A-C and F presented as median ± IQR. Otherwise mean ± SEM.

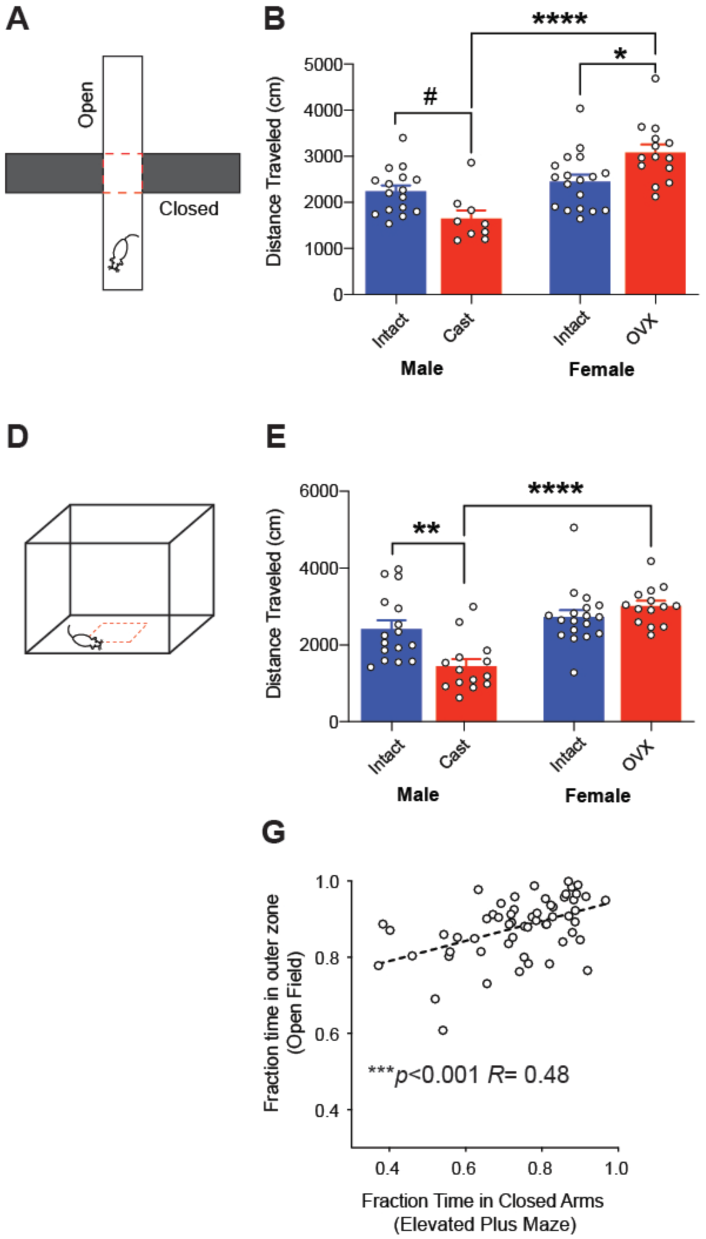


**Figure S2. Divergent effect of prepubertal gonadectomy in males and females on anxiety-like behavior (related to Figure 3).** A. EPM test. B. Total distance traveled during EPM test. OVX females traveled greater distance compared to intact females ($p<0.05$) and Cast males traveled less distance than intact males ($p< 0.05$). OVX females traveled much greater distance compared to Cast males ($p<0.0001$) while there was no difference between intact males and females ($p= 0.32$). C. The fraction time in the closed arms was lower in OVX females compared to intact females ($p< 0.05$) and higher in Cast males compared to intact males ($p< 0.05$). Furthermore, Cast males spent significantly more time in the closed arms compared to OVX females ($p<0.001$). Intact males and females did not differ ($p= 0.77$). D. Open field test. E. Cast males traveled significantly less distance in open field compared to intact males ($p<0.01$). Cast males also traveled significantly less distance compared to OVX females ($p<0.0001$), while intact males and females did not differ in total distance traveled ($p= 0.29$). F. Cast males spent significantly more time in the outer perimeter of the open field compared to intact males ($p<0.001$) as well as OVX females ($p<0.001$) while intact males and females did not differ ($p= 0.08$). All data presented as mean ± SEM, statistical comparisons are Kruskal-Wallis test with post hoc uncorrected Dunn's test (A-E) or one-way ANOVA with uncorrected post hoc unpaired t-test with Welch's correction.

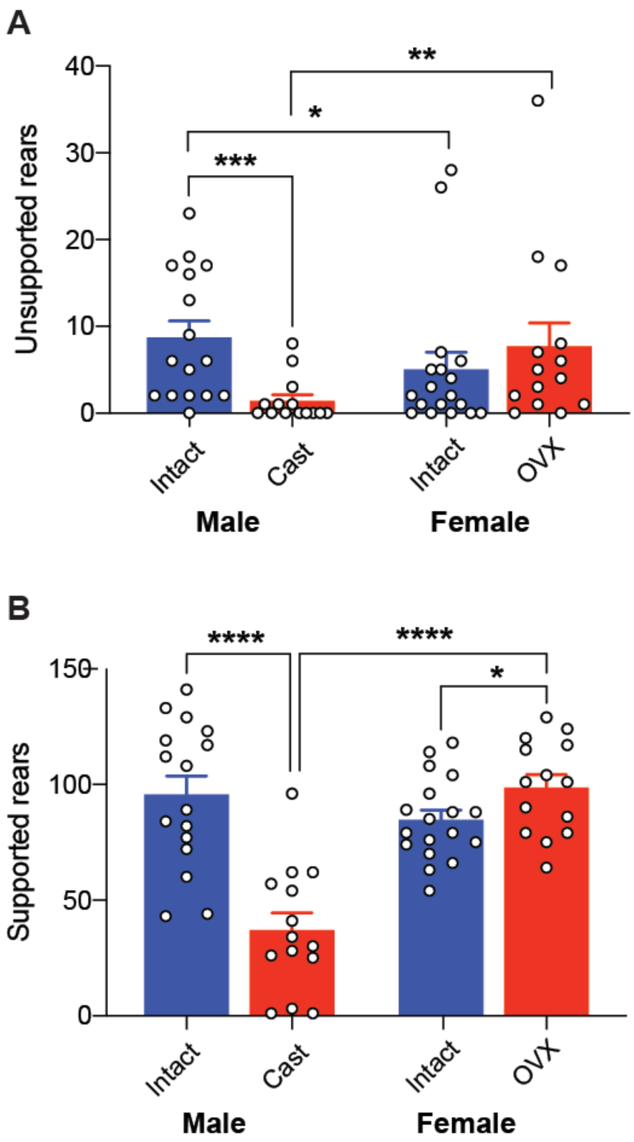


**Figure S3. Divergent effect of prepubertal gonadectomy in males and females on vertical rearing in open field (related to Figure 3).** A. Cast males made fewer unsupported rears compared to intact males (U= 30, uncorrected p= 0.0003) and OVX females (U= 41.5, uncorrected p= 0.0067). Intact males made significantly more unsupported rears than intact females (U= 85.6, uncorrected p= 0.0455), but there was no significant difference between intact females and OVX females (U= 94, uncorrected p= 0.228). B. For supported rears, which occurred along the walls of the arena, Cast males made fewer compared to intact males ($t_{28}$= 5.441, uncorrected p< 0.0001) and OVX females ($t_{26}$= 6.79, uncorrected p<0.0001). There was no significant difference in supported rears between intact males and females ($t_{23.15}$= 1.242, uncorrected p= 0.227), but OVX females made more supported rears compared to intact females ($t_{30}$= 2.08, uncorrected p= 0.046). Data presented as mean ± SEM. *p< 0.05, **p< 0.01, ****p< 0.0001

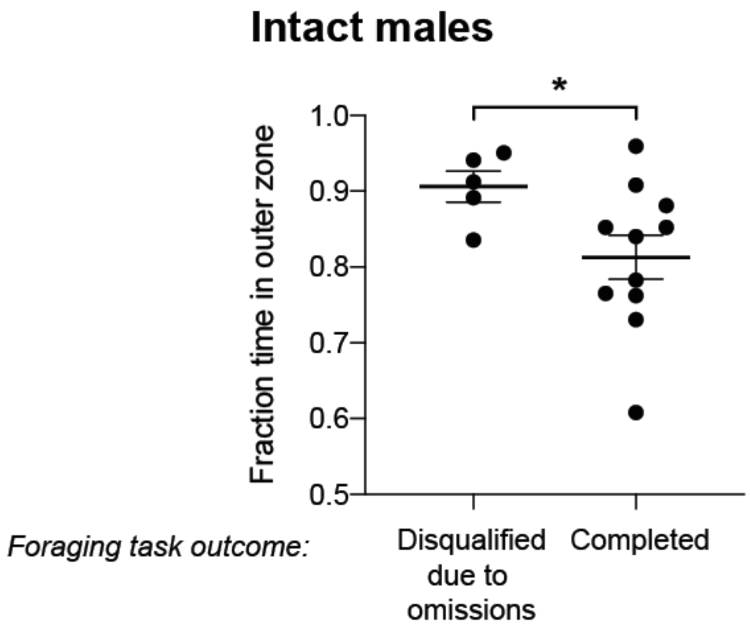


**Figure S4. Intact males show relationship between foraging task completion and (related to Figure 5).** When examined separately, intact males that were disqualified from the foraging task spent more time in the outer fraction of the open field compared to those that successfully completed the reversal phase of the foraging task ($t_{13.86}$= 2.617, p= 0.02 unpaired t test with Welch's correction).
